# Assessing the Clinical Utility of Finite Element Analysis Using Post-operative CT-Derived Models: A Material Comparison of Multi-level Spinal Fusion Constructs

**DOI:** 10.64898/2026.08.05.742715

**Authors:** R Tewari, R.D Johnston, J.M McDonnell, R Storey, S Darwish, J.S Butler, C.M Murphy

**Author notes:** Correspondence to: Dr, Ciara M. Murphy, Department of Anatomy and Regenerative Medicine, Royal College of Surgeons in Ireland, Dublin, Ireland. Both authors contributed equally to this study (R Tewari, R.D Johnston).

## Abstract

Successful instrumented fusion of the lumbar spine is a complex surgical challenge, with positive patient outcomes dependent on careful surgical planning. Material selection is of critical importance to a mechanical construct supporting successful spinal fusion. Therefore, the aims of this study were to (a) evaluate the potential clinical use of finite element analysis (FEA) and (b) conduct a retrospective mechanical analysis of different implant materials in patients having undergone spinal fusion using FEA.

Our methodology involved segmenting the spine from post-operative computed tomography (CT) image data from patients with previous spinal fusion. FEA models representing post-surgery cases were developed and different biomechanical loading conditions such as compression, flexion, bending and extension whilst testing pedicle screws of different materials were simulated.

Patient specific finite element models were created, and biomechanical analysis were completed for all three patients. Polyetheretherketone (PEEK) constructs typically demonstrated lower peak implant stress when compared to titanium constructs for all spinal fusion levels. Furthermore, increasing the spinal fusion level resulted in significant differences in the maximum von Mises stress within both the bone and the instrumentation, whereas the 2-level fusion exhibited comparable stress levels in the bone irrespective of the instrumentation material.

This pilot explores the potential of FEA as a clinical tool for assessing device and bone stresses. In our cohort, different materials can influence the stresses in both the instrumentation and the instrumented vertebrae, suggesting FEA can be useful pre- operative tool with regards to instrument selection and post-operatively to assess instrumentation and bone stresses.

**Graphical Abstract:** 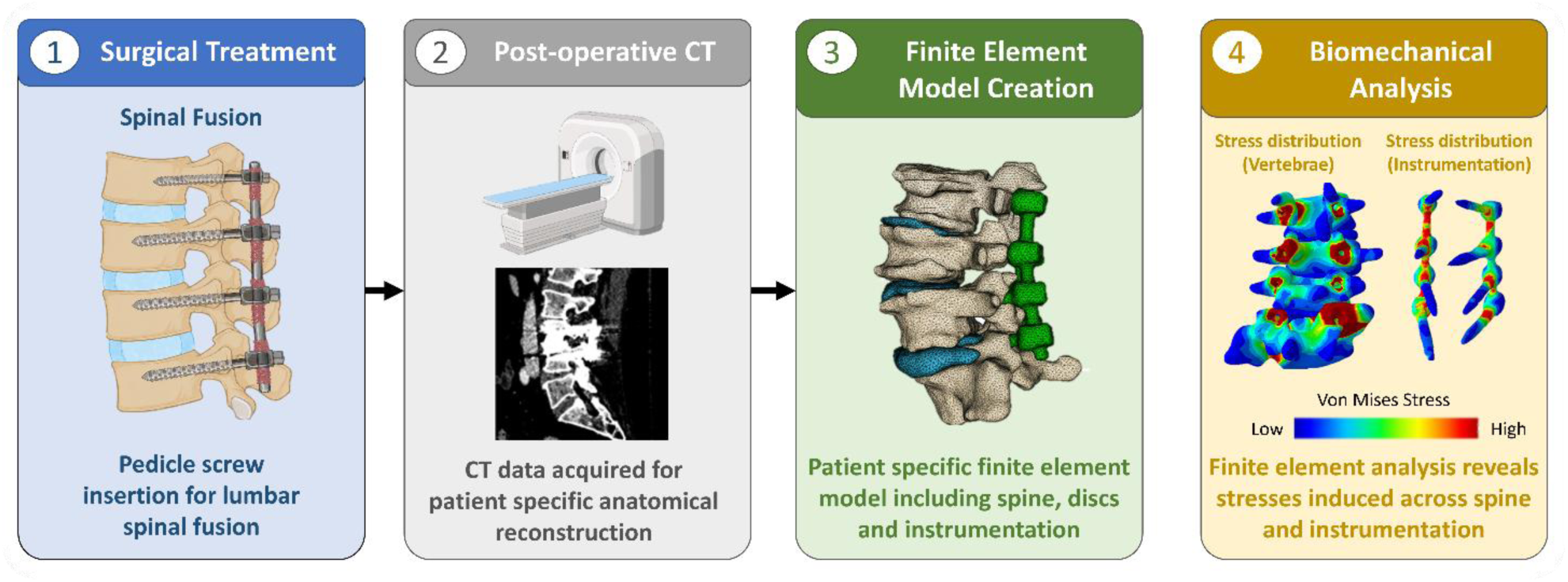

## 1. Background

Lumbar vertebral fractures, often the product of blunt trauma, are a key surgical challenge, with incidence increasing from 14.6 to 22.4 per 100,000 from 2010-2018 in the U.S [1]. These fractures are a key source of morbidity, with patients experiencing pain, decreased mobility and potential neurologic deficit, all of which can substantially impair the quality of life and increase healthcare utilization [2–4]. Furthermore, the burden of spine pathologies is projected to increase with ageing population and rising life expectancy, with degenerative spinal disease and osteoporotic vertebral fractures disproportionally affecting older adults [5,6]. Lumbar fractures are often treated conservatively, typically with physiotherapy, braces or analgesics [7,8]. However, surgical intervention may be indicated in patients with unstable fractures, neurological compromise, progressive deformity, or persistent pain despite conservative management. Surgical treatment typically involves internal spinal fixation, in which bone fragments are repositioned and stabilised using implants such as pedicle screws, or spinal fusion, where biological arthrodesis is achieved to unite two or more vertebrae into one cohesive unit [9–11]. Surgeons must carefully balance several parameters of the given procedure, such as the diameter of the pedicle screws utilised, the positioning of instrumentation and materials utilised [12–16]. Larger and longer screws bestow greater fixation and fusion strength whilst contributing to the distribution of mechanical stress on the vertebrae, however, such screws may have a greater limiting effect on the physiological range of motion and functional outcomes in patients [17,18]. Revision procedures are not uncommon either, with a recent study conducted in the US estimating that the rate of revision surgery for 1 to 2 level lumbar fusion over a 5-year period is 13.5% [19]. Pedicle screw loosening, pullout and instrumentation deformation have been reported in the literature [20–23], with often difficult to treat revision procedures contributing to lost time, greater economic cost and, most importantly, worse patient outcomes. Despite strong advances in intraoperative practices, such as the use of guided navigation and robotics, the positioning and choice of instrumentation is dependent on clinical judgment and experience [24,25]. A prevailing weakness of this approach to pre-operative planning is the lack of patient biomechanical data considered, limiting the degree of patient-specific treatment available [26,27]. Furthermore, surgical teams have poor quantitative methods to determine the risk of instrumentation failure in any given patient.

Finite Element Analysis (FEA) has been traditionally utilised to design instrumentation and assess the stability of orthopaedic implants under various loading conditions [28–30]. Recently, clinicians have taken an interest in the application of FEA to spine surgery, particularly with regards to implant performance and spine stability. Studies have discussed the utilisation of FEA as a pre- and possibly intra-operative planning tool to support patient-specific surgical decision-making [31–34]. Indeed, recent studies have employed FEA pre-operatively in complex tibial or femoral fracture repairs [35,36], with reported reductions in theatre time and length of hospitalisation [36], although a similar application to spine surgery remains limited. Furthermore, an important consideration in spinal fixation is the biomechanical compatibility of implant materials with the native spine. Conventional titanium instrumentation possesses high stiffness and excellent osteointegrative properties, enabling reliable stabilisation and fusion of vertebral segments [37–39]. However, the substantial difference in elastic modulus between titanium and native bone may contribute to stress shielding, altered load transfer, adjacent segment degeneration, and reduced physiological spinal motion [40–42]. In response, alternative biomaterials such as polyether ether ketone (PEEK) have gained increasing interest [43,44]. PEEK is a radiolucent polymer with a lower elastic modulus that more closely approximates cortical bone, potentially allowing for more physiological load sharing and improved biomechanical behaviour following fusion [45–48]. Despite growing interest in PEEK-based instrumentation, its biomechanical performance in multilevel lumbar fusion constructs remains incompletely understood, particularly in the context of patient-specific surgical planning using finite element modelling.

In this study, we aimed to investigate the use of patient-specific finite element analysis (FEA) models generated from postoperative CT imaging to compare the biomechanical performance of different instrumentation strategies for multilevel posterior lumbar fusion. Specifically, we compared the biomechanical behaviour of conventional titanium instrumentation with that of polyetheretherketone (PEEK) constructs under physiologically relevant loading conditions. We hypothesised that the lower stiffness of PEEK instrumentation would promote more physiological stress distribution and load-sharing patterns while maintaining adequate construct stability, supporting its potential as a mechanical alternative to titanium instrumentation in selected spinal fusion procedures.

## 2. Methods

### 2.1 Cohort

This retrospective analysis consisted of a small cohort of patients (N=3) who underwent posterior spinal fusion in a national tertiary referral centre with full ethical approval obtained from the institutional review board. Eligible postoperative CT examinations were selected based on diagnostic image quality, with preference given to datasets exhibiting minimal metal-induced beam-hardening artefacts, complete visualization of the instrumentation and adjacent vertebrae, and no significant image degradation due to patient motion or incomplete acquisition.

The following 3 patients were chosen for this study:

- Patient A underwent L3-L4 posterior instrumented fusion (one motion segment; two instrumented vertebrae).
- Patient B underwent L4-S1 posterior instrumented fusion (two motion segments; three instrumented vertebrae).
- Patient C underwent L2-L5 posterior instrumented fusion (three motion segments; four instrumented vertebrae).

### 2.2 Segmentation

Post-operative radiographic imaging data, specifically Computed Tomography (CT), were obtained for the relevant patients. Segmentations of individual spinal components were created using 3D Slicer (version 5.8.1). The *TotalSegmentor* [49] extension was utilised to create individual masks of the vertebral bodies, which were then manually refined and edited to ensure accuracy due to the substantial metal artefact due to the devices implemented. Intervertebral discs (IVDs) were manually segmented as was the instrumentation. Sufficient quality instrumentation masks were created with thresholding tools and manually refined to remove screw threads. Figure 2 displays the segmented models.

**Figure 1:**
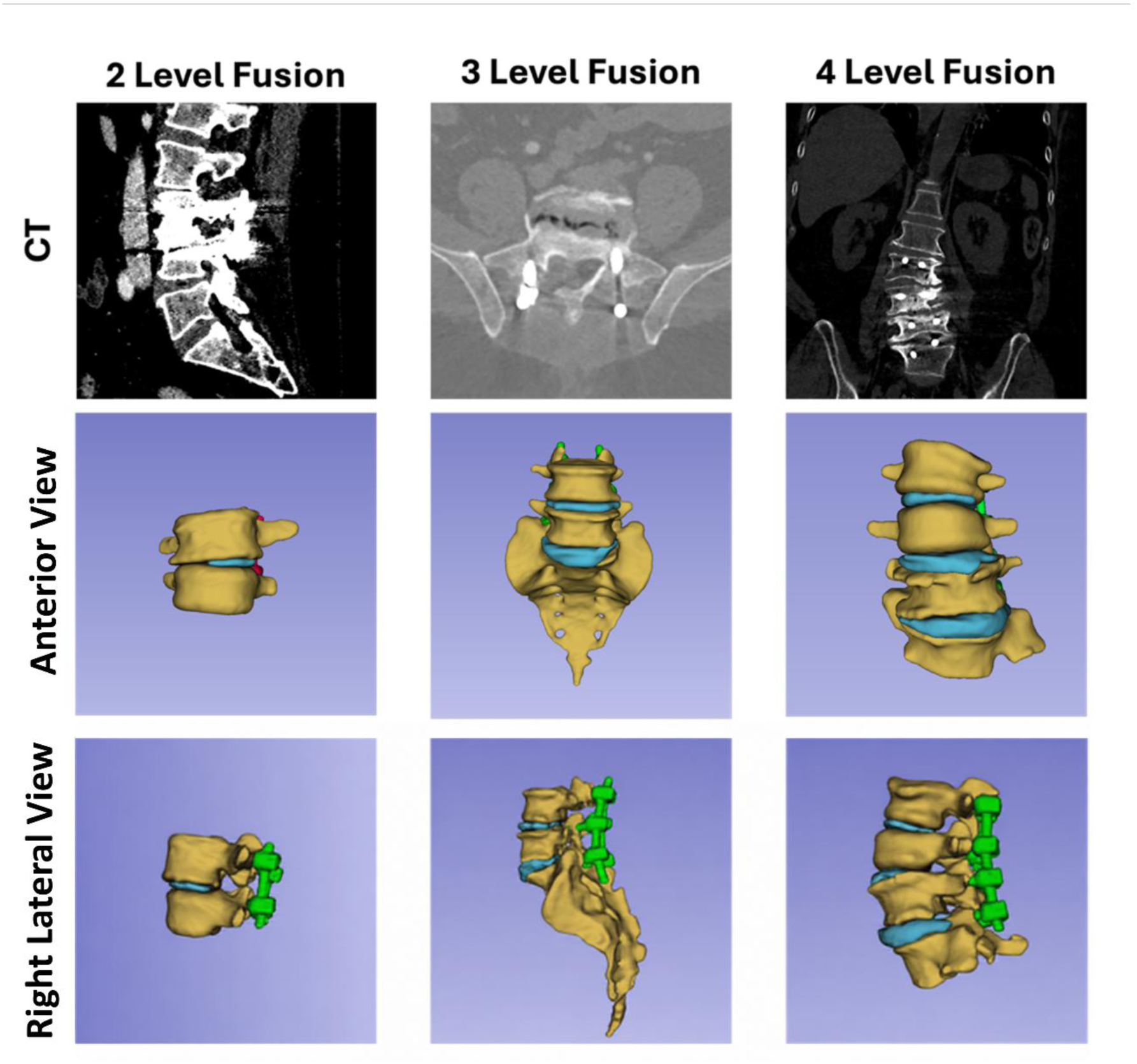
Representative postoperative CT images and corresponding 3D segmentations of 2-level, 3-level, and 4-level spinal fusion cases. The top row shows the CT images, while the middle and bottom rows show anterior and right lateral views of the segmented anatomy, respectively. Vertebrae are shown in yellow, intervertebral discs in blue, and spinal instrumentation (pedicle screws and rods) in green.

**Figure 2:**
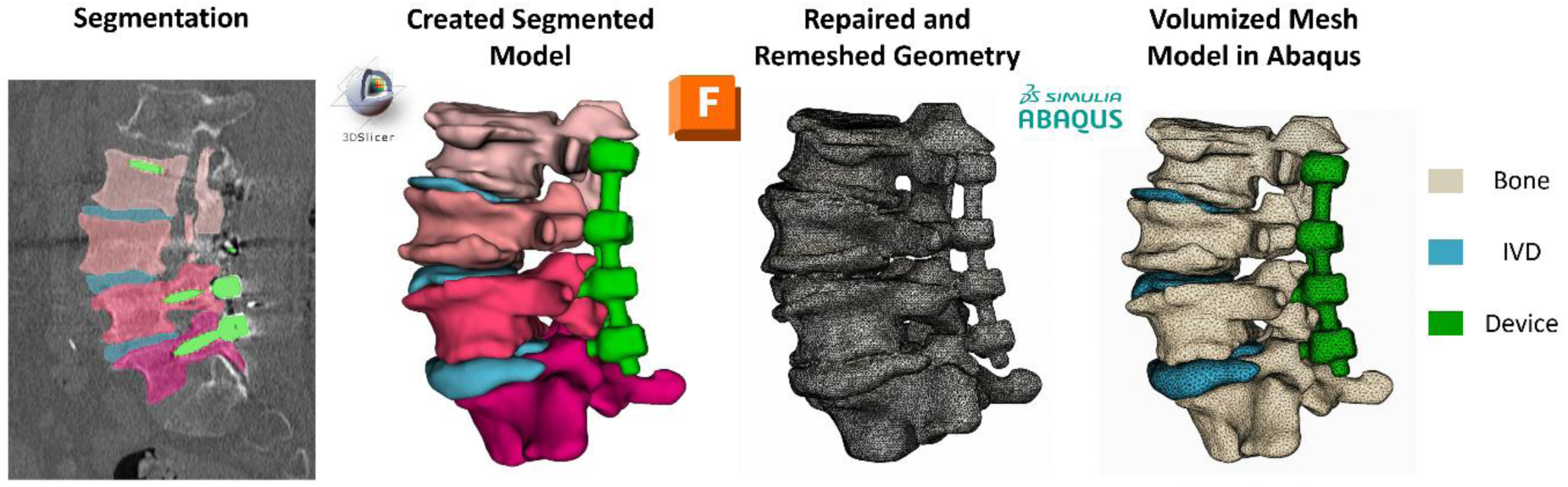
Step by step process to get from post-operative CT imaging to finite element model.

Stereolithography (STL) surface meshes were exported as STL files from the segmentation software and modified using Autodesk Fusion 360 (version 2.0.21550; Autodesk, Inc., San Rafael, CA, USA). Each segmentation was remeshed, repaired and reduced to an appropriate number of faces to preserve geometry whilst reducing computational complexity of the models. Each surface mesh was then volumized and imported into Abaqus (Dassault Systèmes SIMULIA Corp., Johnston, RI, USA, 2023). This process is summarised in Figure 2.

### 2.3 Finite Element Analysis

Patient-specific FE models were created in Abaqus (Dassault Systèmes SIMULIA Corp., Johnston, RI, USA, 2023) for the 2-, 3- and 4-level fusion procedures. Material properties were assigned as per Table 1. Bone and IVDs were modelled as a single homogeneous material to allow for direct comparison between implant materials while limiting model complexity.

**Table 1:** Implemented elastic material parameters for constituents and materials.

| Material | Density (kg/m<sup>3</sup>) | Young's Modulus (MPa) | Poisson's Ratio |
| --- | --- | --- | --- |
| Bone (Homogenous) | 1800 | 12,000 | 0,3 |
| Intervertebral Disc | 1100 | 8 | 0.45 |
| Ti-6Al-4V-ELI | 4430 | 119,000 | 0.34 |
| PEEK | 1300 | 3840 | 0.38 |

Fixed boundary conditions (encastre) were applied in the U1 (radial), U2 (circumferential) and U3 (axial) directions on the caudal vertebra endplate in each fusion model. Loading conditions were applied to the superior endplate of the cranial vertebra. Four loading conditions were simulated: axial compression, flexion, extension, and left coronal-plane lateral bending. A 280 N load was applied to the cranial endplate in each condition. This is summarised in Figure 3. The simulations were conducted using Abaqus/Explicit, ensuring convergence and stability of the solutions. Tetrahedral elements (C3D4) were used to mesh the geometry of the models, due to the irregular nature of these structures. Peak von Mises stress within instrumentation and the adjacent bone was the primary output of interest.

**Figure 3:**
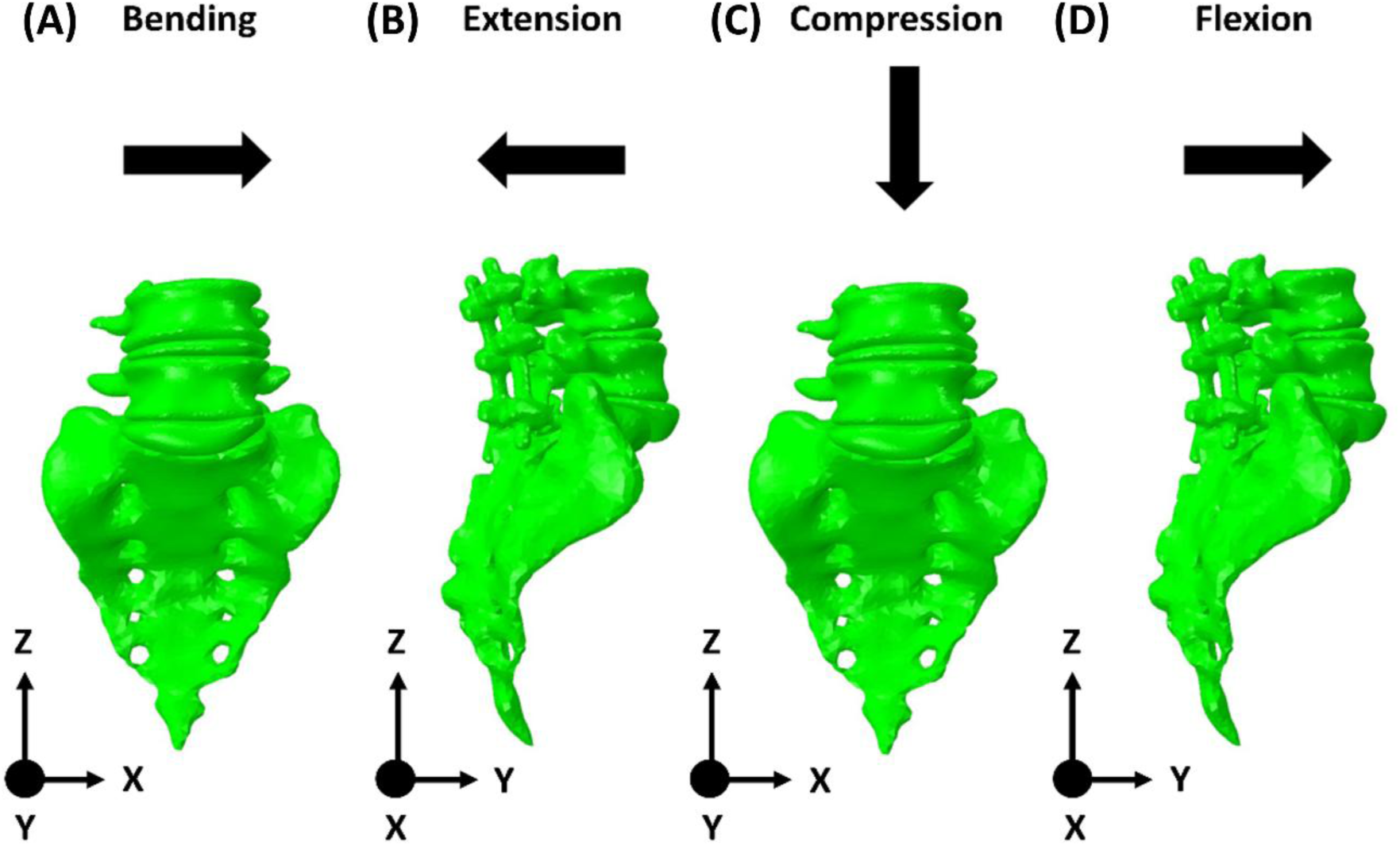
Biomechanical movements simulated for assessment of different levels of spinal fusion (A) Bending (B) Extension (C) Compression (D) Flexion.

Peak Von Mises stress (PVMS) was analysed in 2-, 3-, and 4-level posterior spinal fusion models under flexion, extension, compression and lateral bending (left). Comparisons were performed between titanium and PEEK instrumentation for each construct length and loading condition. To compare materials, we chose a window limit of 100 MPa for figures related to construct alone and a limit of 20 MPa for bone. Further stress distribution analysis was performed by stratifying VMS into five bands: <1 MPa, 1–5 MPa, 5–10 MPa, 10–20 MPa, and >20 MPa. These thresholds were selected to characterise the spectrum of stress magnitudes experienced by cortical and trabecular bone under the applied loading conditions. The lower bands (<1 MPa and 1–5 MPa) represent low physiological stress levels, predominantly associated with trabecular bone [50]. The intermediate bands (5–10 MPa and 10–20 MPa) correspond to regions of functional load transfer within the bone, reflecting increased mechanical demand [51]. The highest band (>20 MPa) identifies localized stress concentrations that may indicate areas of elevated load transmission and potential mechanical overloading.

## 3. Results

### 3.1. 2-Level Fusion

Finite element analysis demonstrated distinct stress distributions between PEEK and titanium instrumentation under bending, extension, compression, and flexion loading conditions, see Figure 4. Across all loading modes, stress concentrations were primarily localised at the vertebral endplates and screw–rod junctions. Under lateral bending, vertebral bone stress was higher in the PEEK (47.75 MPa) compared with the titanium construct (44.56 MPa). Instrumentation stresses were comparatively low and similar between materials, with maximum PVMS values in titanium instrumentation (22.55 MPa) than in the PEEK instrumentation (20.37 MPa), see Figure 4A. During extension, vertebral bone stress was higher in the PEEK construct (62.59 MPa) than in the titanium construct (57.19 MPa). In contrast, instrumentation stress was substantially higher in the titanium instrumentation (48.21 MPa) compared with the PEEK instrumentation (7.87 MPa), see Figure 4B. Compression produced the highest stresses observed for two-level fusion. Vertebral bone stress was marginally higher in the titanium construct (298.6 MPa) than in the PEEK construct (297.8 MPa). However, instrumentation stress was markedly greater in the titanium instrumentation (306.92 MPa) compared with the PEEK instrumentation (98.02 MPa). Stress concentrations under compression were predominantly located around the pedicle screw interfaces and posterior fixation elements, see Figure 4C. Under flexion, vertebral bone stress was higher in the PEEK construct (61.96 MPa) than in the titanium construct (57.22 MPa). Like extension, instrumentation stress was considerably higher in the titanium instrumentation (46.41 MPa) than in the PEEK instrumentation (9.62 MPa), see Figure 4D. All stresses observed for the instrumentation and bone in the 2-level fusion simulations are summarised in Table 2. Overall, vertebral stress patterns were broadly comparable between PEEK and titanium constructs across all loading conditions. However, titanium instrumentation consistently demonstrated greater stress concentration within the fixation hardware, particularly during compression, flexion, and extension, whereas PEEK constructs exhibited lower implant stresses and a more distributed load-sharing pattern.

**Figure 4:**
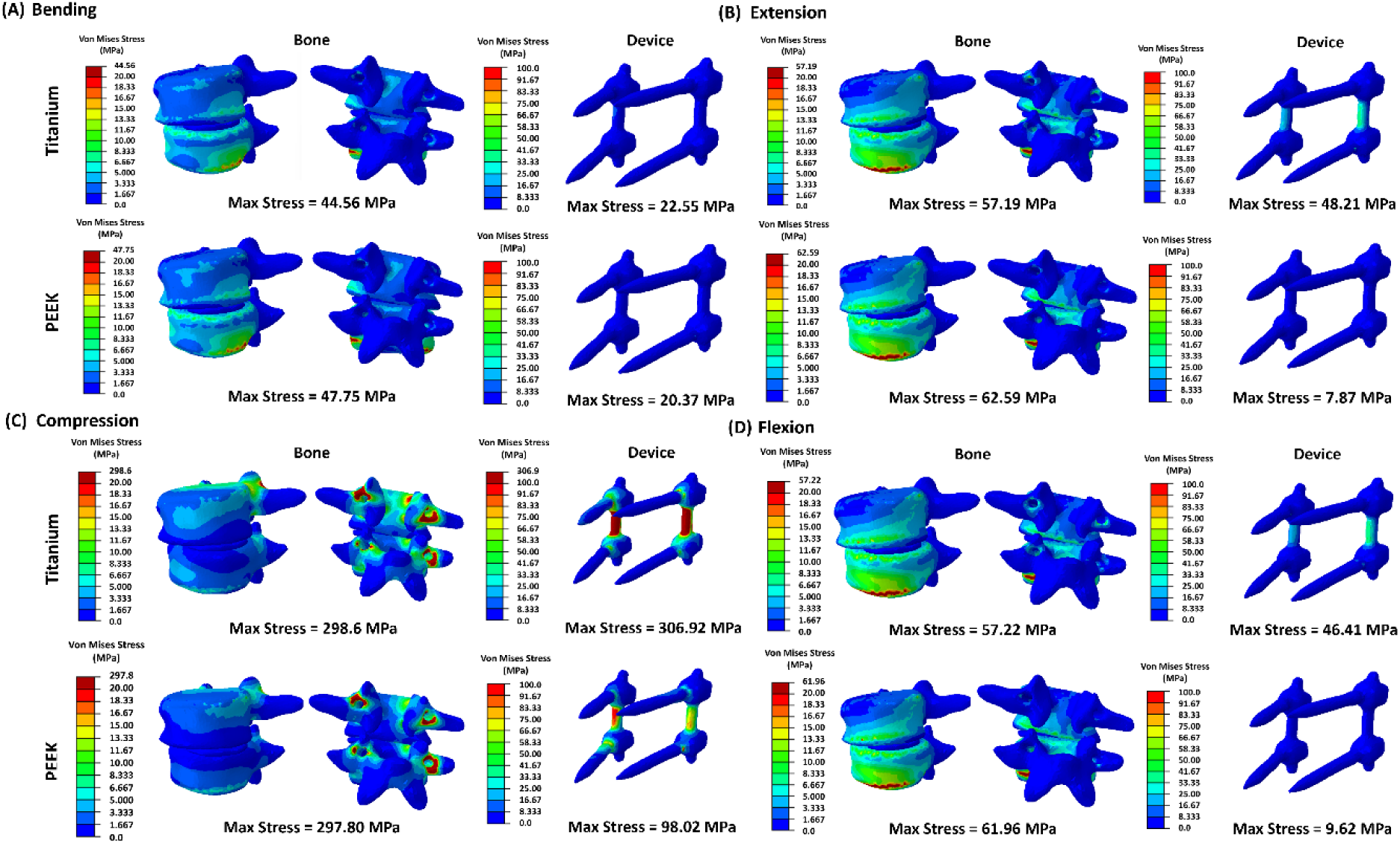
Von Mises stress observed in the bone and instrumentation for both Ti6Al4V and PEEK constructs undergoing (A) Bending (B) Extension (C) Compression (D) Flexion for the 2-level fusion case.

**Table 2:** Summary of stresses observed in instrumentation and vertebrae when either Titanium or PEEK is used for each loading condition for the 2-level fusion case.

| ***Stress in***<br>***Instrumentation***<br>***(MPa)*** | **Bending** | **Extension** | **Compression** | **Extension** |
| --- | --- | --- | --- | --- |
| *Titanium* | 22.55 | 48.21 | 306.92 | 46.41 |
| *PEEK* | 20.37 | 7.87 | 98.02 | 9.62 |

| ***Stress in***<br>***Vertebrae (MPa)*** | **Bending** | **Extension** | **Compression** | **Extension** |
| --- | --- | --- | --- | --- |
| *Titanium* | 44.56 | 57.19 | 298.60 | 57.22 |
| *PEEK* | 47.75 | 62.59 | 297.80 | 61.96 |

### 3.2. 3-Level Fusion

Simulation of the 3-level lumbosacral fixation constructs also demonstrated distinct stress distributions between titanium and PEEK instrumentation under bending, extension, compression, and flexion loading conditions, see Figure 5. Under lateral bending, vertebral bone stress was marginally higher in the PEEK construct (29.67 MPa) than in the titanium construct (29.21 MPa). Instrumentation stresses as substantially higher in the titanium instrumentation (21.91 MPa) compared with the PEEK instrumentation (1.97 MPa), see Figure 5A. During extension, vertebral bone stress was higher in the titanium construct (88.41 MPa) than in the PEEK construct (53.97 MPa). Similarly, Instrumentation stresses also differed considerably between materials, with considerably higher stresses within the titanium instrumentation (55.63 MPa) compared with the PEEK instrumentation (12.96 MPa). Stress concentrations were primarily observed at the superior screw–rod junctions and adjacent vertebral endplates, see Figure 5B. Compression generated the highest stress magnitudes across all loading conditions. Vertebral bone stress was higher in the titanium construct (341.82 MPa) than in the PEEK construct (297.80 MPa). The most pronounced divergence occurred within the instrumentation, where instrumentation stress was markedly higher in the titanium construct (790.24 MPa) than in the PEEK construct (98.02 MPa). High stress concentrations were localised around the pedicle screw necks and rod junctions, indicating substantial load transfer through the titanium fixation system, see Figure 5C. Under flexion, vertebral bone stress was marginally higher in the titanium construct (55.75 MPa) than in the PEEK construct (54.77 MPa). However, titanium instrumentation again demonstrated greater stress amplification, with instrumentation stress was again higher in the titanium instrumentation (46.41 MPa) compared with the PEEK instrumentation (20.93 MPa), see Figure 5D. All stresses observed for the instrumentation and bone in the 3-level fusion simulations are summarised in Table 3. Overall, vertebral stress distributions varied modestly between the two materials, whereas instrumentation stresses were consistently higher in the titanium constructs. The greatest differences were observed under compression and extension loading, where titanium fixation demonstrated marked stress concentration within the hardware, while PEEK constructs exhibited lower implant stresses and a more distributed load-sharing behaviour.

**Figure 5:**
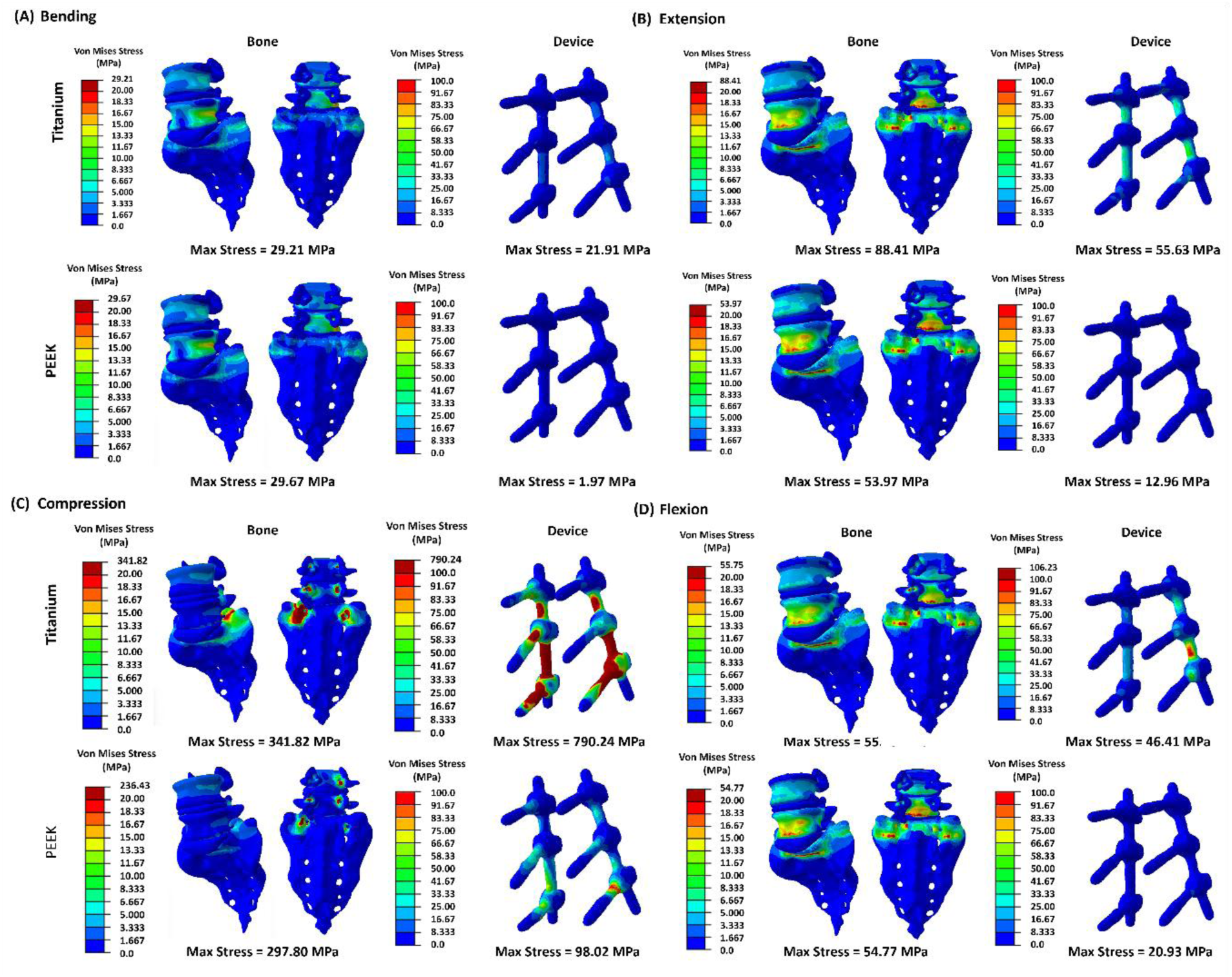
Von Mises stress observed in the bone and instrumentation for both Ti6Al4V and PEEK undergoing (A) Bending (B) Extension (C) Compression (D) Flexion for the 3-level fusion case.

**Table 3:** Summary of stresses observed in instrumentation and vertebrae when either Titanium or PEEK is used for each loading condition for the 3-level fusion case.

| ***Stress in***<br>***Instrumentation***<br>***(MPa)*** | **Bending** | **Extension** | **Compression** | **Extension** |
| --- | --- | --- | --- | --- |
| *Titanium* | 21.91 | 55.63 | 790.24 | 46.41 |
| *PEEK* | 1.97 | 12.96 | 98.02 | 20.93 |

| ***Stress in***<br>***Vertebrae (MPa)*** | **Bending** | **Extension** | **Compression** | **Extension** |
| --- | --- | --- | --- | --- |
| *Titanium* | 29.21 | 88.41 | 341.82 | 55.75 |
| *PEEK* | 29.67 | 53.97 | 297.80 | 54.77 |

### 3.3. 4-Level Fusion

The most pronounced differences in material performance with marked differences in stress distributions being observed in the 4-level constructs, see Figure 6. Under lateral bending, vertebral bone stress was substantially higher in the titanium construct (144.68 MPa) than in the PEEK construct (46.86 MPa). Instrumentation stress was also markedly higher in the titanium instrumentation (660.67 MPa) compared with the PEEK instrumentation (17.07 MPa). Pronounced stress concentration was observed along the posterior rod system in the titanium construct, see Figure 6A. During extension, vertebral bone stress was higher in the PEEK construct (75.89 MPa) than in the titanium construct (42.43 MPa). Despite the lower vertebral bone stress, instrumentation stress remained considerably higher in the titanium instrumentation (155.69 MPa) compared with the PEEK instrumentation (27.82 MPa). Stress concentrations were predominantly located around the superior fixation segments and rod curvature regions, see Figure 6B. Compression generated the highest instrumentation stresses overall. Vertebral bone stress was marginally higher in the titanium construct (117.54 MPa) than in the PEEK construct (114.82 MPa). However, instrumentation stress was substantially higher in the titanium instrumentation (850.16 MPa) compared with the PEEK instrumentation (86.53 MPa). These high stresses were concentrated at the screw necks and along the connecting rods, indicating significant load transfer through the metallic fixation system, see Figure 6C. Under flexion, vertebral bone stress was higher in the titanium construct (69.21 MPa) than in the PEEK construct (43.01 MPa). Instrumentation stress was again markedly higher in the titanium instrumentation (248.66 MPa) compared with the PEEK instrumentation (35.85 MPa), see Figure 6D. All stresses observed for the instrumentation and bone in the 3-level fusion simulations are summarised in Table 4.

**Figure 6:**
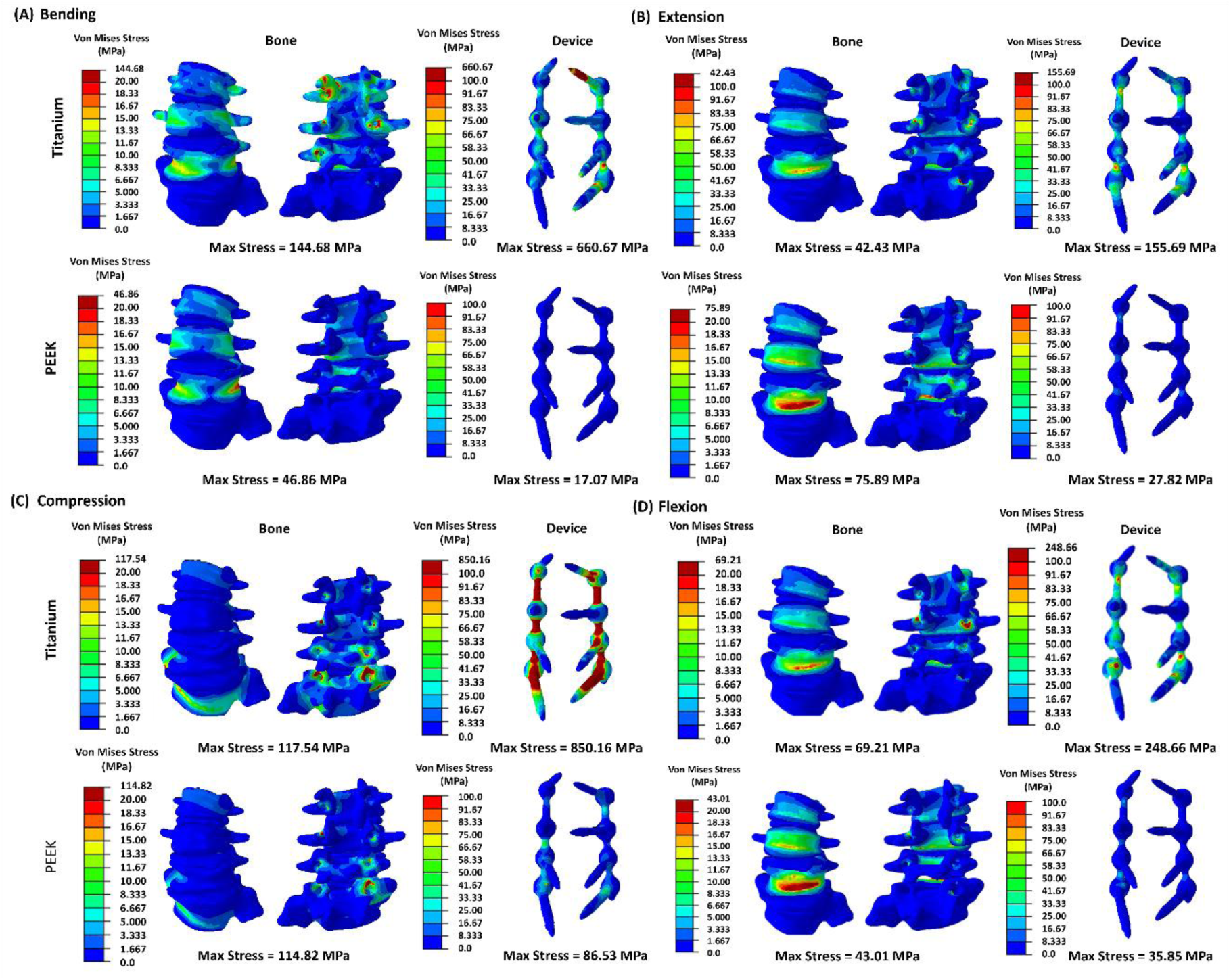
Von Mises stress observed in the bone and instrumentation for both Ti6Al4V and PEEK instrumentation undergoing (A) Bending (B) Extension (C) Compression (D) Flexion for the 4-level fusion case.

**Table 4:**
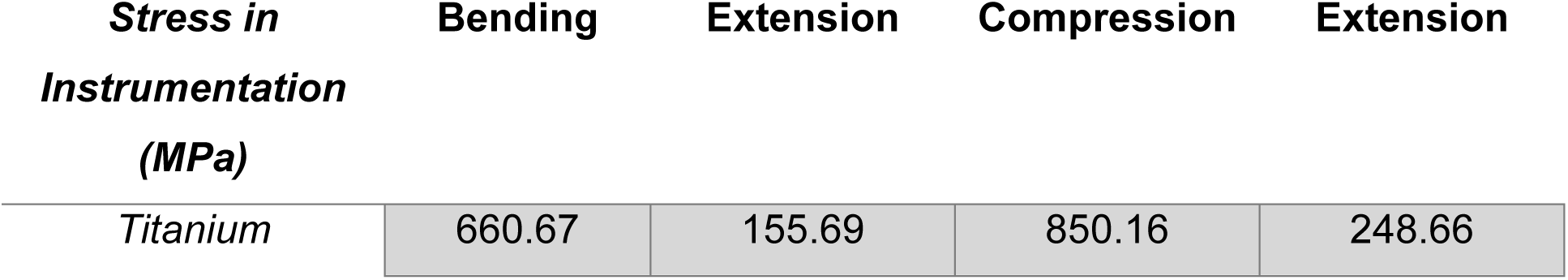
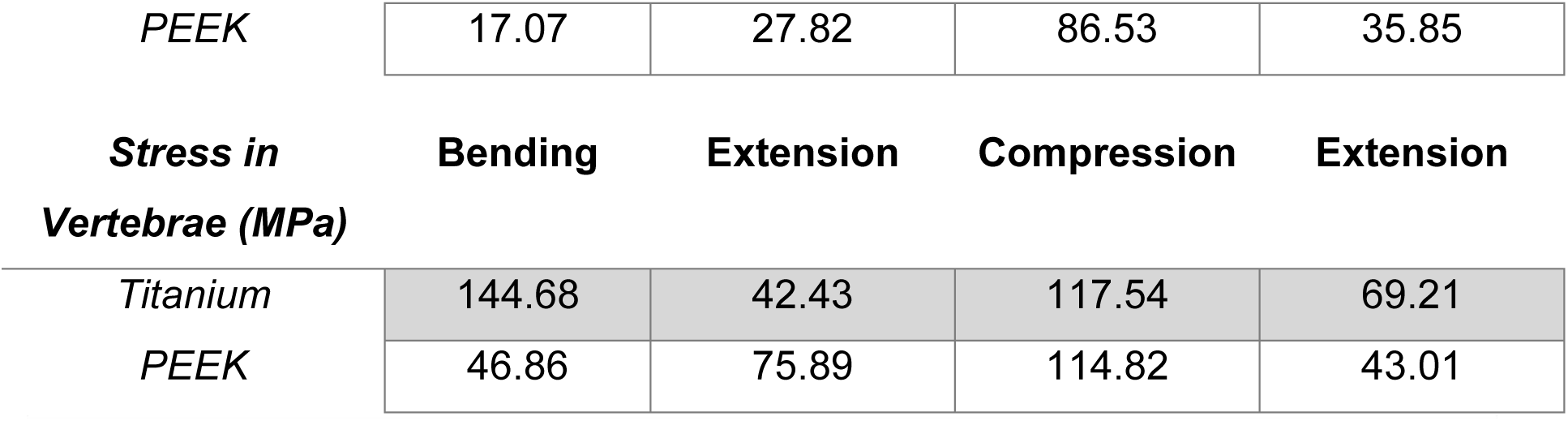
Summary of stresses observed in instrumentation and vertebrae when either Titanium or PEEK is used for each loading condition for the 2-level fusion case.

### 3.4. Percentage volume analysis across spinal fusion levels

Analysis of vertebral stress distribution demonstrated that most of the vertebral volume across all constructs and loading conditions experienced low stress magnitudes (<5 MPa). However, notable differences were observed between construct materials and fixation lengths with respect to the proportion of vertebral volume exposed to higher stress bands.

Under lateral bending, the greatest proportion of vertebral volume was concentrated within the <1 MPa and 1–5 MPa stress bands for all constructs. Three-level constructs, particularly the 3-level PEEK model, demonstrated the highest percentage of vertebral volume within the <1 MPa range, exceeding 60% of total vertebral volume. In contrast, 2-level constructs exhibited a greater proportion of tissue within the 1–5 MPa stress band. Only a small proportion of vertebral volume (<10%) experienced stresses between 5–10 MPa, while stresses exceeding 10 MPa were negligible across all models, see Figure 7A. During extension, a similar distribution pattern was observed, with most vertebral volume remaining within the <1 MPa and 1–5 MPa ranges. Three-level constructs again demonstrated the largest proportion of low-stress vertebral volume, whereas 2-level constructs showed a relative shift toward the 1–5 MPa and 5–10 MPa bands. Titanium and PEEK constructs demonstrated broadly similar distributions, although PEEK models generally showed slightly greater proportions within the lowest stress range, see Figure 7B. Compression loading produced the greatest redistribution of vertebral stress volumes. Across all constructs, the proportion of vertebral volume within the <1 MPa range increased substantially, particularly in the 3-level PEEK model, which demonstrated the highest low-stress volume overall. However, 2-level constructs exhibited a marked increase in vertebral volume within the 1–5 MPa and 5–10 MPa stress bands, indicating greater stress transfer to the surrounding vertebrae. Small but detectable proportions of vertebral volume also extended into the 10–20 MPa and >20 MPa bands, particularly in the titanium constructs, see Figure 7C. Under flexion, stress distributions were again predominantly concentrated below 5 MPa. Four-level and three-level constructs demonstrated larger proportions of vertebral volume within the <1 MPa range, whereas 2-level constructs demonstrated greater volumes within the 1–5 MPa and 5– 10 MPa bands. Minimal vertebral volume exceeded 10 MPa across all configurations, see Figure 7D. Overall, longer fixation constructs (3- and 4-level) demonstrated a greater proportion of vertebral volume within lower stress bands, suggesting more distributed load-sharing across the vertebral column. In contrast, shorter 2-level constructs showed increased vertebral volume exposed to moderate stress ranges, particularly during compression and flexion. Differences between titanium and PEEK constructs were modest overall; however, PEEK models generally demonstrated slightly greater proportions of low-stress vertebral volume, consistent with a more evenly distributed stress profile

**Figure 7:**
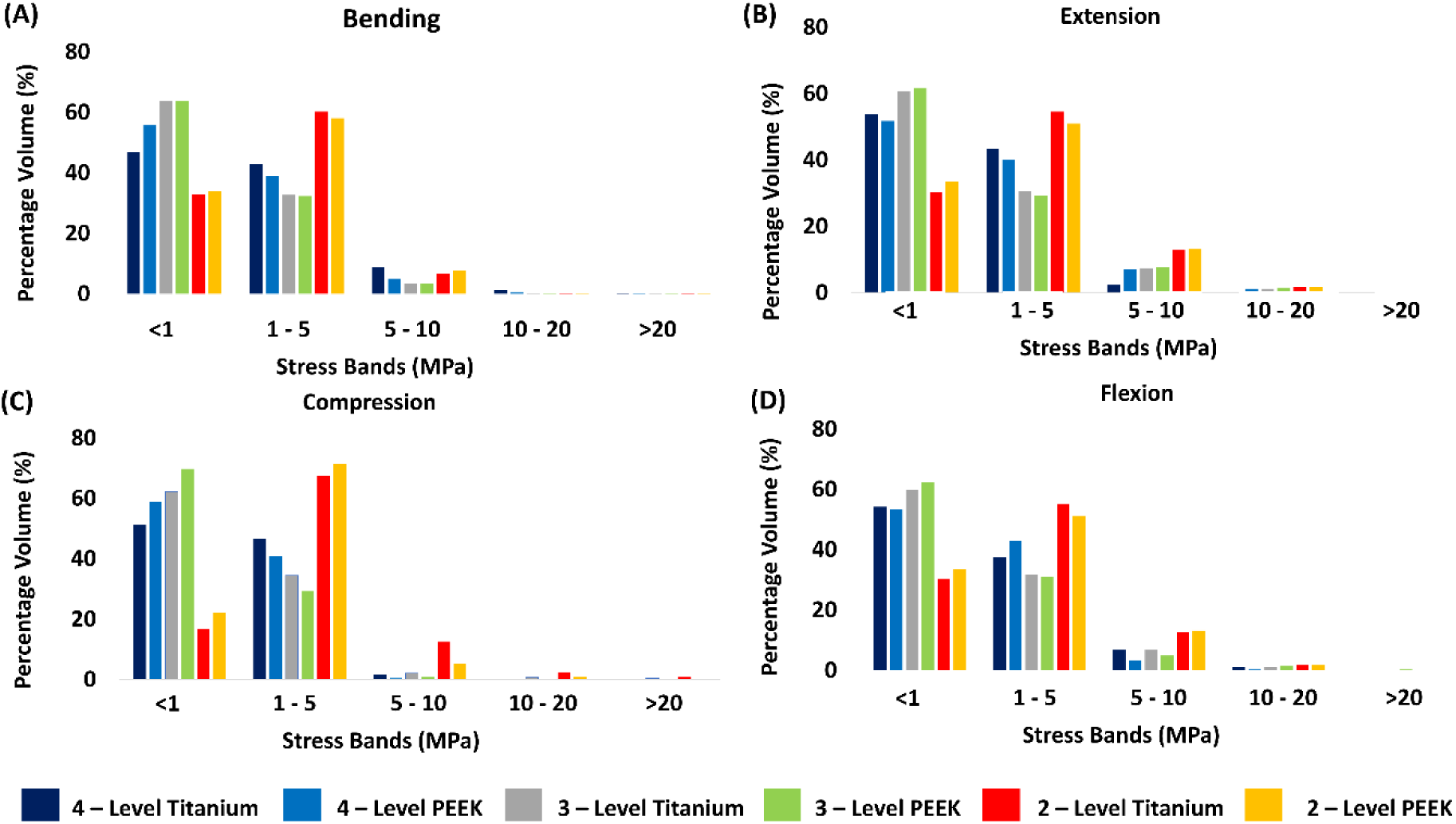
Percentage volume of elements experiencing stresses under (A) Bending (B) Extension (C) Compression and (D) Flexion loading for Titanium and PEEK constructs incorporated into 2-Level, 3-Level and 4-Level fusion simulations.

## 4. Discussion

Lumbar fusion is performed at high volume internationally, with trends suggesting increasing use of multilevel constructs [52,53]. Failed procedures, particularly in the context of instrumentation failure, remain a potential adverse consequence of surgical intervention with subsequent complex revision surgery required. FEA has been proposed as a method to better optimise pre-operative planning and to achieve greater personalised surgery. This study investigates the technical feasibility of using finite element analysis (FEA) to evaluate stress distributions in bone and spinal instrumentation by comparing the biomechanical performance of different instrumentation materials in 2-, 3-, and 4-level posterior spinal fusion constructs.

In our study, we have demonstrated that both construct length and implant material influence the stress distribution within the vertebrae and the fixation hardware. Firstly, across all loading conditions simulated, titanium instrumentation consistently exhibited higher peak von Mises stresses within the screws and rods, whereas PEEK constructs demonstrated lower implant stresses and more distributed load sharing pattern. These findings are consistent with previous FEA investigations showing that PEEK based fixation systems reduce stress concentrations within instrumentation due to their lower elastic modulus and similarity to cortical bone [54–56]. The most pronounced differences observed in our simulations were under compression loading, where titanium constructs generated markedly greater stresses within the instrumentation compared with PEEK, with similar trends also observed during flexion and extension. This behaviour is likely related to the increased stiffness of titanium, which concentrates load within the fixation hardware, whereas the more compliant PEEK material allows greater force transmission to the surrounding vertebral structures. Previous studies similarly reported that titanium rods absorb a larger proportion of applied load, while PEEK promotes more physiological load sharing and reduces implant stress amplification [57–59]. However, these biomechanical advantages should be interpreted with caution. While the lower stiffness of PEEK may reduce implant stress concentrations and potentially minimise stress shielding, it may also permit greater construct deformation and intersegmental micromotion [60]. Although a degree of micromotion may promote physiological load transfer, excessive motion at the fusion site could compromise the mechanical environment required for successful treatment [61]. Consequently, the clinical implications of reduced construct stiffness likely represent a trade-off between decreasing implant stress and maintaining sufficient stability to support fusion. Despite the reduction in implant stress observed with PEEK instrumentation, vertebral stresses remained broadly comparable between materials in most loading conditions. This suggests that PEEK constructs may maintain construct stability within the constraints of this model while reducing stress concentrations within the hardware itself. Similar observations have been reported in biomechanical analyses comparing CFR-PEEK and titanium systems, where reduced implant stresses occurred without substantial increases in vertebral loading [54,62].

Secondly, fixation length also influenced the stress distribution throughout the vertebral column. Three- and four-level fusions demonstrated a larger proportion of vertebral volume within the lowest stress bands (<1 MPa), suggesting that loads were distributed more evenly across multiple spinal segments. In contrast, two-level constructs exhibited greater proportions of vertebral volume within the 1–5 MPa and 5–10 MPa stress bands, particularly during compression and flexion. These findings indicate that shorter fusions may concentrate stress over fewer motion segments, potentially increasing the mechanical burden on adjacent vertebrae and terminal fixation points. From a clinical perspective, the elevated implant stresses observed in titanium constructs may contribute to implant fatigue, rod fracture, and screw loosening over time. Furthermore, the high stiffness of titanium has been associated with stress shielding, whereby reduced physiological loading of the surrounding bone may contribute to bone resorption and adjacent segment degeneration. In contrast, the lower stiffness of PEEK allows more physiological force transmission through the vertebral column, may reduce stress-shielding effects, although this requires validation against fusion, loosening, fatigue, and clinical outcome data.

Overall, the present findings suggest that implant material has a greater influence on instrumentation stress than on vertebral stress, with PEEK constructs consistently reducing stress concentrations within the fixation system. Increasing construct length further promoted distributed load sharing throughout the vertebral column. Together, these findings support the potential biomechanical advantages of PEEK instrumentation in reducing hardware stress concentrations while maintaining a more physiological mechanical environment. However, biological considerations and intended outcomes should be considered in terms in any device / material selection [63,64].

### 4.1. Clinical Feasibility

Another objective of this study was to explore the potential of finite element analysis (FEA) as a pre- and postoperative clinical decision-support tool. Our findings suggest that FEA may ultimately allow clinicians to quantify the biomechanical consequences of different surgical interventions using anatomically accurate, patient-specific digital models of the spine. This capability could support more informed surgical planning and enable treatment strategies to be better tailored to the anatomical and biomechanical characteristics of individual patients. However, clear hurdles still remain. For example, even if this technology were fully validated, it would not immediately fit into clinical workflows. Segmentations, even with the approach detailed in the methods and practice, may take several hours to produce depending on the quality of the CT imaging. Issues with image quality such as the beam hardening effect evident in Figure 1, also necessitate the need for pre-operative CT and possibly magnetic resonance imaging, (MRI) which patients may not be indicated for. The setup of each model in Abaqus is time consuming. The main difficulty arose when remeshing the components, as this was found to be the most difficult and cumbersome aspect of the entire procedure and often involved a degree of trial and error as well. Although, once setup was achieved, it was straightforward to test different loading conditions and materials. In terms of the running the model, these would often take an hour or more to calculate depending on the complexity of the model. Workflow automation, validated segmentation pipelines, and clinically interpretable thresholds will be required before routine clinical deployment. Lastly, for FEA to be useful in routine clinical practice, its outputs must be presented in terms that directly inform surgical decision-making rather than as complex stress distributions alone. Clinicians are more likely to benefit from concise, actionable information, such as identification of vertebral levels at increased risk of screw failure, estimates of overall construct stiffness, recommendations for reinforcement or augmentation strategies where appropriate, and an indication of the confidence or uncertainty associated with these predictions. Presenting results in this manner would improve interpretability and facilitate integration into surgical planning.

### 4.2. Limitations and Future Work

This comparative study is an exploratory application of FEA in the context of pre- operative surgical decision making. As such there are several limitations which must be acknowledged. Firstly, bone was modelled as a single homogeneous material for simplicity, without accounting for the distinct biomechanical properties of cortical and cancellous bone that can be derived from CT-based material mapping [65]. Accurate conversion of Hounsfield units to bone elastic modulus could not be performed from the post-operative CT images due to severe beam-hardening artefacts caused by the instrumentation. Future studies should incorporate pre-operative CT imaging to enable patient-specific assignment of material properties and provide a more robust biomechanical assessment. Secondly, IVDs were also modelled as a single homogenous material, which comprises the annulus fibrosus, a highly anisotropic tissue, and the nucleus pulposus, which behaves more isotropically. Complementary MRI would allow for segmentation of the components to be included into the FEA model more readily and should be included in future work. In addition, the models did not include spinal ligaments, facet joints, paraspinal muscles, physiological follower loads and the screw-bone interaction was simplified. Furthermore, the models were not validated against cadaveric or benchtop testing, nor were the predictions correlated with clinical outcomes. Lastly, incorporating a greater number of patients would also be appropriate in future studies. This would allow assessment of inter-patient variability in spinal anatomy, bone quality, and loading characteristics, all of which may substantially influence construct biomechanics and stress distribution. Inclusion of patients with varying degrees of bone density, deformity severity, and fusion length would help establish whether the observed trends between PEEK and titanium instrumentation are consistent across different clinical scenarios.

## 5. Conclusions

To conclude, this study provides proof-of-concept evidence supporting the potential of finite element analysis (FEA) as a clinically relevant decision-support tool. Within our cohort, implant material influenced stress distributions according to both fusion construct and patient-specific anatomy, suggesting that FEA could assist preoperative planning by informing implant and material selection. Furthermore, this study establishes a computational framework for evaluating the biomechanical performance of different fixation constructs and investigating factors associated with implant success or failure.

## 6. Ethical Approval

Ethical approval was acquired from the Mater Hospital ethics committee for the obtaining of images used in this study.

## References

[1] J. Wakim, T. Rajan, A. Beschloss, A. Albayar, A. Ozturk, C. Saifi, Etiologies, incidence, and demographics of lumbar vertebral fractures in U.S. emergency departments, J. Spine Surg. 8 (2022) 21–28. 10.21037/jss-21-110.

[2] S. Rajasekaran, R.M. Kanna, A.P. Shetty, Management of thoracolumbar spine trauma An overview, Indian J. Orthop. 49 (2015) 72–82. 10.4103/0019-5413.143914.

[3] G. Vilà-Canet, A. García de Frutos, A. Covaro, M.T. Ubierna, E. Caceres, Thoracolumbar fractures without neurological impairment: A review of diagnosis and treatment, EFORT Open Rev. 1 (2016) 332–338. 10.1302/2058-5241.1.000029.

[4] T. Liu, H. Zou, H. Zou, Global Burden of Vertebral Fractures and Spinal Cord Injuries Due to Falls From 1990 to 2021: A Population-Based Study, Glob. Spine J. 15 (2025) 3743–3755. 10.1177/21925682251331442.

[5] J.L. Dieleman, J. Cao, A. Chapin, C. Chen, Z. Li, A. Liu, C. Horst, A. Kaldjian, T. Matyasz, K.W. Scott, A.L. Bui, M. Campbell, H.C. Duber, A.C. Dunn, A.D. Flaxman, C. Fitzmaurice, M. Naghavi, N. Sadat, P. Shieh, E. Squires, K. Yeung, C.J.L. Murray, US Health Care Spending by Payer and Health Condition, 1996-2016, JAMA - J. Am. Med. Assoc. 323 (2020) 863–884. 10.1001/jama.2020.0734.

[6] M.L. Ferreira, K. De Luca, L.M. Haile, J.D. Steinmetz, G.T. Culbreth, M. Cross, J.A. Kopec, P.H. Ferreira, F.M. Blyth, R. Buchbinder, J. Hartvigsen, A.M. Wu, S. Safiri, A.D. Woolf, G.S. Collins, K.L. Ong, S.E. Vollset, A.E. Smith, J.A. Cruz, K.G. Fukutaki, S.M. Abate, M. Abbasifard, M. Abbasi-Kangevari, Z. Abbasi-Kangevari, A. Abdelalim, A. Abedi, H. Abidi, Q.E.S. Adnani, A. Ahmadi, R.O. Akinyemi, A.T. Alamer, A.Z. Alem, Y. Alimohamadi, M.A. Alshehri, M.M. Alshehri, H. Alzahrani, S. Amini, S. Amiri, H. Amu, C.L. Andrei, T. Andrei, B. Antony, J. Arabloo, J. Arulappan, A. Arumugam, T. Ashraf, S.S. Athari, N. Awoke, S. Azadnajafabad, T.W. Bärnighausen, L.H. Barrero, A. Barrow, A. Barzegar, L.M. Bearne, I.M. Bensenor, A.Y. Berhie, B.B. Bhandari, V.S. Bhojaraja, A. Bijani, B.B.A. Bodicha, S.R. Bolla, J. Brazo-Sayavera, A.M. Briggs, C. Cao, P. Charalampous, V.K. Chattu, F.M. Cicuttini, B. Clarsen, S. Cuschieri, O. Dadras, X. Dai, L. Dandona, R. Dandona, A. Dehghan, T.G.G. Demie, E. Denova-Gutiérrez, S.M.R. Dewan, S.D. Dharmaratne, M.L. Dhimal, M. Dhimal, D. Diaz, M. Didehdar, L.E. Digesa, M. Diress, H.T. Do, L.P. Doan, M. Ekholuenetale, M. Elhadi, S. Eskandarieh, S. Faghani, J. Fares, A. Fatehizadeh, G. Fetensa, I. Filip, F. Fischer, R.C. Franklin, B. Ganesan, B.N.B. Gemeda, M.E. Getachew, A. Ghashghaee, T.K. Gill, M. Golechha, P. Goleij, B. Gupta, N. Hafezi-Nejad, A. Haj-Mirzaian, P.K. Hamal, A. Hanif, N.I. Harlianto, H. Hasani, S.I. Hay, J.J. Hebert, G. Heidari, M. Heidari, R. Heidari-Soureshjani, M.M. Hlongwa, M.S. Hosseini, A.K. Hsiao, I. Iavicoli, S.E. Ibitoye, I.M. Ilic, M.D. Ilic, S.M.S. Islam, M.D. Janodia, R.P. Jha, H.A. Jindal, J.B. Jonas, G.G. Kabito, H. Kandel, R.J. Kaur, V.R. Keshri, Y.S. Khader, E.A. Khan, M.J. Khan, M.A.B. Khan, H.R.K. Kashani, J. Khubchandani, Y.J. Kim, A. Kisa, J. Klugarová, A.A. Kolahi, H.R. Koohestani, A. Koyanagi, G.A. Kumar, N. Kumar, T. Lallukka, S. Lasrado, W.C. Lee, Y.H. Lee, A. Mahmoodpoor, J.N. Malagón-Rojas, M.R. Malekpour, R. Malekzadeh, N. Malih, M.M. Mehndiratta, E.M. Nasab, R.G. Menezes, A.F.A. Mentis, M.K. Mesregah, T.R. Miller, M. Mirza-Aghazadeh-Attari, M. Mobarakabadi, Y. Mohammad, E. Mohammadi, S. Mohammed, A.H. Mokdad, S. Momtazmanesh, L. Monasta, M.A. Moni, E. Mostafavi, C.J.L. Murray, T.S. Nair, J. Nazari, S.A. Nejadghaderi, S. Neupane, S.N. Kandel, C.T. Nguyen, A. Nowroozi, H. Okati-Aliabad, E. Omer, A. Oulhaj, M.O. Owolabi, S. Panda-Jonas, A. Pandey, E.K. Park, S. Pawar, P. Pedersini, J. Pereira, M.F.P. Peres, I.R. Petcu, M. Pourahmadi, A. Radfar, S. Rahimi- Dehgolan, V. Rahimi-Movaghar, M. Rahman, A.M. Rahmani, N. Rajai, C.R. Rao, V. Rashedi, M.M. Rashidi, Z.A. Ratan, D.L. Rawaf, S. Rawaf, A.M.N. Renzaho, N. Rezaei, Z. Rezaei, L. Roever, G. De Andrade Ruela, B. Saddik, A. Sahebkar, S. Salehi, F. Sanmarchi, S.G. Sepanlou, S. Shahabi, S. Shahrokhi, E. Shaker, M.B. Shamsi, M. Shannawaz, S. Sharma, M. Shaygan, R.A. Sheikhi, J.K. Shetty, R. Shiri, S. Shivalli, P. Shobeiri, M.M. Sibhat, A. Singh, J.A. Singh, H. Slater, M. Solmi, R. Somayaji, K.K. Tan, R. Thapar, S.A. Tohidast, S.V. Tahbaz, R. Valizadeh, T.J. Vasankari, N. Venketasubramanian, V. Vlassov, B. Vo, Y.P. Wang, T. Wiangkham, L. Yadav, A. Yadollahpour, S.H.Y. Jabbari, L. Yang, F. Yazdanpanah, N. Yonemoto, M.Z. Younis, I. Zare, A. Zarrintan, M. Zoladl, T. Vos, L.M. March, Global, regional, and national burden of low back pain, 1990–2020, its attributable risk factors, and projections to 2050: a systematic analysis of the Global Burden of Disease Study 2021, Lancet Rheumatol. 5 (2023) e316–e329. 10.1016/S2665-9913(23)00098-X.

[7] I.K. Genev, M.K. Tobin, S.P. Zaidi, S.R. Khan, F.M.L. Amirouche, A.I. Mehta, Spinal Compression Fracture Management: A Review of Current Treatment Strategies and Possible Future Avenues., Glob. Spine J. 7 (2017) 71–82. 10.1055/s-0036-1583288.

[8] M. Squires, J.H. Green, R. Patel, I. Aleem, Clinical outcomes after bracing for vertebral compression fractures: a systematic review and meta-analysis of randomized trials, J. Spine Surg. 9 (2023) 139–148. 10.21037/jss-22-78.

[9] A.R. Vaccaro, G.D. Schroeder, C.K. Kepler, F. Cumhur Oner, L.R. Vialle, F. Kandziora, J.D. Koerner, M.F. Kurd, M. Reinhold, K.J. Schnake, J. Chapman, A. Aarabi, M.G. Fehlings, M.F. Dvorak, The surgical algorithm for the AOSpine thoracolumbar spine injury classification system, Eur. Spine J. 25 (2016) 1087–1094. 10.1007/s00586-015-3982-2.

[10] C. Court, C. Vincent, Percutaneous fixation of thoracolumbar fractures: Current concepts, Orthop. Traumatol. Surg. Res. 98 (2012) 900–909. 10.1016/j.otsr.2012.09.014.

[11] M. Bydon, R. De La Garza-Ramos, M. Macki, A. Baker, A.K. Gokaslan, A. Bydon, Lumbar fusion versus nonoperative management for treatment of discogenic low back pain: A systematic review and meta-analysis of randomized controlled trials, J. Spinal Disord. Tech. 27 (2014) 297–304. 10.1097/BSD.0000000000000072.

[12] G.F. Solitro, K. Whitlock, F. Amirouche, A.I. Mehta, A. McDonnell, Currently adopted criteria for pedicle screw diameter selection, Int. J. Spine Surg. 13 (2019) 132–145. 10.14444/6018.

[13] W. Cho, S.K. Cho, C. Wu, The biomechanics of pedicle screw-based instrumentation, J. Bone Jt. Surg. - Ser. B 92 (2010) 1061–1065. 10.1302/0301-620X.92B8.24237.

[14] K. Phan, J. Hogan, M. Maharaj, R.J. Mobbs, Cortical Bone Trajectory for Lumbar Pedicle Screw Placement: A Review of Published Reports, Orthop. Surg. 7 (2015) 213–221. 10.1111/os.12185.

[15] C. Tsagkaris, A.K. Calek, M.R. Fasser, J.M. Spirig, S. Caprara, M. Farshad, J. Widmer, Bone density optimized pedicle screw insertion, Front. Bioeng. Biotechnol. 11 (2023) 1–8. 10.3389/fbioe.2023.1270522.

[16] A. Warburton, S.J. Girdler, C.M. Mikhail, A. Ahn, S.K. Cho, Biomaterials in spinal implants: A review, Neurospine 17 (2020) 101–110. 10.14245/ns.1938296.148.

[17] K. Matsukawa, T. Konomi, K. Matsubayashi, J. Yamane, Y. Yato, Influence of Pedicle Screw Insertion Depth on Posterior Lumbar Interbody Fusion: Radiological Significance of Deeper Screw Placement, Glob. Spine J. 14 (2024) 470–477. 10.1177/21925682221110142.

[18] K. Matsukawa, Y. Yato, H. Imabayashi, Impact of Screw Diameter and Length on Pedicle Screw Fixation Strength in Osteoporotic Vertebrae:A Finite Element Analysis, Asian Spine J. 15 (2021) 566–574. 10.31616/asj.2020.0353.

[19] D. Cummins, K. Hindoyan, H.H. Wu, A.A. Theologis, M. Callahan, B. Tay, S. Berven, Reoperation and Mortality Rates Following Elective 1 to 2 Level Lumbar Fusion: A Large State Database Analysis, Glob. Spine J. 12 (2022) 1708–1714. 10.1177/2192568220986148.

[20] D.P. Falk, R. Agrawal, B. Dehghani, R. Bhan, S. Gupta, M.C. Gupta, Instrumentation Failure in Adult Spinal Deformity Patients, J. Clin. Med. 13 (2024) 1–18. 10.3390/jcm13154326.

[21] S. Sumiya, K. Fukushima, Y. Kurosa, T. Hirai, H. Inose, T. Yoshii, A. Okawa, Comparative analysis of clinical factors associated with pedicle screw pull-out during or immediately after surgery between intraoperative cone-beam computed tomography and postoperative computed tomography, BMC Musculoskelet. Disord. 22 (2021) 1–10. 10.1186/s12891-020-03916-9.

[22] V. Krishnan, V. Varghese, G.S. Kumar, N. Yoganandan, Identification of pedicle screw pullout load paths for osteoporotic vertebrae, Asian Spine J. 14 (2020) 273–279. 10.31616/ASJ.2019.0174.

[23] L. Marie-Hardy, H. Pascal-Moussellard, A. Barnaba, R. Bonaccorsi, C. Scemama, Screw Loosening in Posterior Spine Fusion: Prevalence and Risk Factors, Glob. Spine J. 10 (2020) 598–602. 10.1177/2192568219864341.

[24] C. McNamee, D. Kelly, J.M. McDonnell, A.M. Sowa, A. Menon, S. Darwish, J.S. Butler, The learning curve of robotic assisted pedicle screw placement: individual patient data meta-analysis, Spine J. 26 (2026) 180–188. 10.1016/j.spinee.2025.07.007.

[25] J. Maalouly, M. Sarkar, J. Choi, Retrospective study assessing the learning curve and the accuracy of minimally invasive robot-assisted pedicle screw placement during the first 41 robot-assisted spinal fusion surgeries, Mini-Invasive Surg. 5 (2021). 10.20517/2574-1225.2021.57.

[26] J. Berhouet, R. Samargandi, Emerging Innovations in Preoperative Planning and Motion Analysis in Orthopedic Surgery, Diagnostics 14 (2024). 10.3390/diagnostics14131321.

[27] F. Galbusera, G. Casaroli, T. Bassani, Artificial intelligence and machine learning in spine research, JOR Spine 2 (2019) 1–20. 10.1002/jsp2.1044.

[28] D.A. Sin, D.H. Heo, Comparative finite element analysis of lumbar cortical screws and pedicle screws in transforaminal and posterior lumbar interbody fusion, Neurospine 16 (2019) 298–304. 10.14245/ns.1836030.015.

[29] M. Xu, J. Yang, I.H. Lieberman, R. Haddas, Finite element method-based study of pedicle screw–bone connection in pullout test and physiological spinal loads, Med. Eng. Phys. 67 (2019) 11–21. 10.1016/j.medengphy.2019.03.004.

[30] C.J. Kim, S.M. Son, S.H. Choiw, T.S. Goh, J.S. Lee, C.S. Lee, Numerical evaluation of spinal stability after posterior spinal fusion with various fixation segments and screw types in patients with osteoporotic thoracolumbar burst fracture using finite element analysis, Appl. Sci. 11 (2021). 10.3390/app11073243.

[31] M. Ahmadi, H. Chen, M. Lin, D. Biswas, J. Doulgeris, Y. Tang, E.D. Engeberg, J. Hashemi, G. Pires, F.D. Vrionis, Streamlined and efficient patient-specific modeling for lumbar spine segmentation and finite element analysis, Sci. Rep. 15 (2025) 1–23. 10.1038/s41598-025-19664-6.

[32] A. Verma, A. Jain, S. Sekhar Sethy, V. Verma, N. Goyal, M. Vathulya, P. Kandwal, Finite element analysis and its application in Orthopaedics: A narrative review, J. Clin. Orthop. Trauma 58 (2024) 102803. 10.1016/j.jcot.2024.102803.

[33] R. Wang, Z. Wu, Recent advancement in finite element analysis of spinal interbody cages: A review, Front. Bioeng. Biotechnol. 11 (2023) 1–17. 10.3389/fbioe.2023.1041973.

[34] M.C. Wang, A. Kiapour, E. Massaad, J.H. Shin, N. Yoganandan, A guide to finite element analysis models of the spine for clinicians, J. Neurosurg. Spine 40 (2024) 38–44. 10.3171/2023.7.SPINE23164.38.

[35] Y. He, Y. Liu, B. Yin, D. Wang, H. Wang, P. Yao, J. Zhou, Application of Finite Element Analysis Combined With Virtual Computer in Preoperative Planning of Distal Femoral Fracture, Front. Surg. 9 (2022) 1–9. 10.3389/fsurg.2022.803541.

[36] Z. Xu, Y. Li, S. Tian, X. Xu, H. Zhou, M. Yang, Preliminary exploration of finite element biomechanical preoperative planning for complex tibial plateau fractures, Sci. Rep. 15 (2025) 1–15. 10.1038/s41598-025-01085-0.

[37] F.A. Shah, P. Thomsen, A. Palmquist, Osseointegration and current interpretations of the bone-implant interface, Acta Biomater. 84 (2019) 1–15. 10.1016/j.actbio.2018.11.018.

[38] A.J. Croft, H. Chanbour, J.W. Chen, M.W. Young, B.F. Stephens, Implant Surface Technologies to Promote Spinal Fusion: A Narrative Review, Int. J. Spine Surg. 17 (2023) S35–S43. 10.14444/8559.

[39] K.A.K. Raju, A. Biswas, Surface modifications and coatings to improve osseointegration and antimicrobial activity on titanium surfaces: A statistical review over the last decade, J. Orthop. 67 (2025) 68–87. 10.1016/j.jor.2025.01.002.

[40] J. Li, Z. Du, S. Cao, T. Lu, Z. Sun, H. Wei, H. Li, T. Zhang, Quantitative relationships between elastic modulus of rod and biomechanical properties of transforaminal lumbar interbody fusion: a finite element analysis, Front. Bioeng. Biotechnol. 12 (2024) 1–9. 10.3389/fbioe.2024.1510597.

[41] K. Li, S. Cao, J. Chen, J. Qin, B. Yuan, J. Li, Determining a relative total lumbar range of motion to alleviate adjacent segment degeneration after transforaminal lumbar interbody fusion: a finite element analysis, BMC Musculoskelet. Disord. 25 (2024) 1–9. 10.1186/s12891-024-07322-3.

[42] J. Litak, M. Szymoniuk, W. CzyZewski, Z. Hoffman, J. Litak, L. Sakwa, P. Kamieniak, Metallic Implants Used in Lumbar Interbody Fusion, Materials (Basel). 15 (2022) 1–23. 10.3390/ma15103650.

[43] S.M. Kurtz, J.N. Devine, PEEK biomaterials in trauma, orthopedic, and spinal implants, Biomaterials 28 (2007) 4845–4869. 10.1016/j.biomaterials.2007.07.013.

[44] Z. Wei, Z. Zhang, W. Zhu, X. Weng, Polyetheretherketone development in bone tissue engineering and orthopedic surgery, Front. Bioeng. Biotechnol. 11 (2023) 1–16. 10.3389/fbioe.2023.1207277.

[45] Y.C. Yao, P.H. Chou, H.H. Lin, S.T. Wang, M.C. Chang, Outcome of Ti/PEEK Versus PEEK Cages in Minimally Invasive Transforaminal Lumbar Interbody Fusion, Glob. Spine J. 13 (2023) 472–478. 10.1177/21925682211000323.

[46] A. Chahlavi, Reduced Subsidence With PEEK-Titanium Composite Versus 3D Titanium Cages in a Retrospective, Self-Controlled Study in Transforaminal Lumbar Interbody Fusion, Glob. Spine J. 15 (2025) 1598–1607. 10.1177/21925682241253168.

[47] W. Huang, W. Wang, X. Xu, L. Wang, J. Wang, X. Yu, Radiological outcomes of PEEK rods in patients with lumbar degenerative diseases: A minimum 5- year follow-up, Front. Surg. 10 (2023) 1–7. 10.3389/fsurg.2023.1146893.

[48] A.G. Anghel, J. Garthmann, B. Alkahawagi, A PEEK-Based Pedicle Screw System for One-Level Lumbar Spinal Canal Stenosis: An Appraisal at a Five-Year Follow Up, J. Clin. Med. 14 (2025) 3–5. 10.3390/jcm14124252.

[49] J. Wasserthal, H.C. Breit, M.T. Meyer, M. Pradella, D. Hinck, A.W. Sauter, T. Heye, D.T. Boll, J. Cyriac, S. Yang, M. Bach, M. Segeroth, TotalSegmentator: Robust Segmentation of 104 Anatomic Structures in CT Images, Radiol. Artif. Intell. 5 (2023). 10.1148/ryai.230024.

[50] T.M. Keaveny, E.F. Morgan, G.L. Niebur, O.C. Yeh, BIOMECHANICS OF TRABECULAR BONE, Annu. Rev. Biomed. Eng. (2001) 307–33.

[51] S. Eswaran, A. Gupta, M. Adams, T. Keaveny, Cortical and Trabecular Load Sharing in the Human Vertebral Body, J. Bone Miner. Res. 21 (2006) 193–206. 10.1359/JBMR.051027.

[52] M.J. Reisener, M. Pumberger, J. Shue, F.P. Girardi, A.P. Hughes, Trends in lumbar spinal fusion—a literature review, J. Spine Surg. 6 (2020) 752–76. 10.21037/jss-20-492.

[53] B.I. Martin, S.K. Mirza, B.A. Karamian, A.J. Schoenfeld, H. Ko, P. Suri, D.S. Brodke, Cost and Utilization Trends of Lumbar Fusion, JAMA Netw. Open 9 (2026) 1–13. 10.1001/jamanetworkopen.2026.0452.

[54] S. Oikonomidis, J. Greven, J. Bredow, M. Eh, A. Prescher, H. Fischer, J. Thüring, P. Eysel, F. Hildebrand, P. Kobbe, M.J. Scheyerer, C. Herren, Biomechanical effects of posterior pedicle screw-based instrumentation using titanium versus carbon fiber reinforced PEEK in an osteoporotic spine human cadaver model, Clin. Biomech. 80 (2020) 105153. 10.1016/j.clinbiomech.2020.105153.

[55] C. Li, Y. Zhao, L. Qi, B. Xu, L. Yue, R. Zhu, C. Li, Comparison of biomechanical effects of polyetheretherketone (PEEK) rods and titanium rods in lumbar long-segment instrumentation: a finite element study, Front. Bioeng. Biotechnol. 12 (2024) 1–12. 10.3389/fbioe.2024.1416046.

[56] Y.N. Wang, Y.N. Ren, J. Han, C. Chen, X. Sun, M.Y. Di, Y.M. Dou, X.L. Ma, Z. Wang, C.F. Du, Q. Yang, Biomechanical effects of screws of different materials on vertebra-pediculoplasty: a finite element study, Front. Bioeng. Biotechnol. 11 (2023) 1–7. 10.3389/fbioe.2023.1225925.

[57] J. Li, S. Cao, B. Zhao, Biomechanical comparison of polyetheretherketone rods and titanium alloy rods in transforaminal lumbar interbody fusion: a finite element analysis, BMC Surg. 24 (2024) 1–13. 10.1186/s12893-024-02462-8.

[58] J. Wu, L. Shi, D. Liu, Z. Wu, P. Gao, W. Liu, X. Li, Z. Guo, Evaluating Screw Stability After Pedicle Screw Fixation With PEEK Rods, Glob. Spine J. 13 (2023) 393–399. 10.1177/2192568221996692.

[59] Y.H. Ahn, W.M. Chen, K.Y. Lee, K.W. Park, S.J. Lee, Comparison of the load-sharing characteristics between pedicle-based dynamic and rigid rod devices, Biomed. Mater. 3 (2008). 10.1088/1748-6041/3/4/044101.

[60] D.K. Sengupta, B. Bucklen, P.C. McAfee, J. Nichols, R. Angara, S. Khalil, The Comprehensive Biomechanics and Load-Sharing of Semirigid PEEK and Semirigid Posterior Dynamic Stabilization Systems, Adv. Orthop. 2013 (2013) 1–9. 10.1155/2013/745610.

[61] C.M. Bono, A. Khandha, S. Vadapalli, S. Holekamp, V.K. Goel, S.R. Garfin, Residual sagittal motion after lumbar fusion: A finite element analysis with implications on radiographic flexion-extension criteria, Spine (Phila. Pa. 1976). 32 (2007) 417–422. 10.1097/01.brs.0000255201.74795.20.

[62] R.A. Lindtner, R. Schmid, T. Nydegger, M. Konschake, W. Schmoelz, Pedicle screw anchorage of carbon fiber-reinforced PEEK screws under cyclic loading, Eur. Spine J. 27 (2018) 1775–1784. 10.1007/s00586-018-5538-8.

[63] E. Massaad, N. Fatima, A. Kiapour, M. Hadzipasic, G.M. Shankar, J.H. Shin, Polyetheretherketone versus titanium cages for posterior lumbar interbody fusion: Meta-analysis and review of the literature, Neurospine 17 (2020) 473. 10.14245/ns.2040058.029.c2.

[64] N.A. Patel, S. O’bryant, C.D. Rogers, S. Chakravarti, J. Gendreau, N.J. Brown, Z.A. Pennington, N.B. Hatcher, L.D. Diaz-Aguilar, M.H. Pham, Three-Dimensional-Printed Titanium Versus Polyetheretherketone Cages for Lumbar Interbody Fusion: A Systematic Review of Comparative In Vitro, Animal, and Human Studies, Neurospine 20 (2023) 451–463. 10.14245/ns.2346244.122.

[65] G. Chen, B. Schmutz, D. Epari, K. Rathnayaka, S. Ibrahim, M.A. Schuetz, M.J. Pearcy, A new approach for assigning bone material properties from CT images into finite element models, J. Biomech. 43 (2010) 1011–1015. 10.1016/j.jbiomech.2009.10.040.

